# Novel SARS-CoV-2 Whole-genome sequencing technique using Reverse Complement PCR enables easy, fast and accurate outbreak analysis in hospital and community settings

**DOI:** 10.1101/2020.10.29.360578

**Authors:** Femke Wolters, Jordy P.M. Coolen, Alma Tostmann, Lenneke F.J. van Groningen, Chantal P. Bleeker-Rovers, Edward C.T.H. Tan, Nannet van der Geest-Blankert, Jeannine L.A. Hautvast, Joost Hopman, Heiman F.L. Wertheim, Janette C. Rahamat-Langendoen, Marko Storch, Willem J.G. Melchers

## Abstract

**Background:** Current transmission rates of severe acute respiratory syndrome coronavirus 2 (SARS-CoV-2) are still increasing and many countries are facing second waves of infections. Rapid SARS-CoV-2 whole-genome sequencing (WGS) is often unavailable but could support public health organizations and hospitals in monitoring and determining transmission links. Here we report the use of reverse complement polymerase chain reaction (RC-PCR), a novel technology for WGS of SARS-CoV-2 enabling library preparation in a single PCR saving time, resources and enables high throughput screening. Additionally, we show SARS-CoV-2 diversity and possible transmission within the Radboud university medical center (Radboudumc) during September 2020 using RC-PCR WGS.

**Methods:** A total of 173 samples tested positive for SARS-CoV-2 between March and September 2020 were selected for whole-genome sequencing. Ct values of the samples ranged from 16 to 42. They were collected from 83 healthcare workers and three patients at the Radboudumc, in addition to 64 people living in the area around the hospital and tested by the local health services. For validation purposes, nineteen of the included samples were previously sequenced using Oxford Nanopore Technologies and compared to RC-PCR WGS results. The applicability of RC-PCR WGS in outbreak analysis for public health service and hospitals was tested on six suspected clusters containing samples of healthcare workers and patients with an epidemiological link.

**Findings:** RC-PCR resulted in sequencing data for 146 samples. It showed a genome coverage of up to 98,2% for samples with a maximum Ct value of 32. Comparison to Oxford Nanopore technologies gives a near-perfect agreement on 95% of the samples (18 out of 19). Three out of six clusters with a suspected epidemiological link were fully confirmed, in the others, four healthcare workers were not associated. In the public health service samples, a previously unknown chain of transmission was confirmed.

**Significance statement:** SAR-CoV-2 whole-genome sequencing using RC-PCR is a reliable technique and applicable for use in outbreak analysis and surveillance. Its ease of use, high-trough screening capacity and wide applicability makes it a valuable addition or replacement during this ongoing SARS-CoV-2 pandemic.

**Funding:** None

**Research in context:** *Evidence before this study:* At present whole genome sequencing techniques for SARS-CoV-2 have a large turnover time and are not widely available. Only a few laboratories are currently able to perform large scale SARS-CoV-2 sequencing. This restricts the use of sequencing to aid hospital and community infection prevention.

*Added value of this study:* Here we present clinical and technical data on a novel Whole Genome Sequencing technology, implementing reverse-complement PCR. It is able to obtain high genome coverage of SARS-CoV-2 and confirm and exclude epidemiological links in 173 healthcare workers and patients. The RC-PCR technology simplifies the workflow thereby reducing hands on time. It combines targeted PCR and sequence library construction in a single PCR, which normally takes several steps. Additionally, this technology can be used in concordance with the widely available range of Illumina sequencers.

*Implications of all the available evidence:* RC-PCR whole genome sequencing technology enables rapid and targeted surveillance and response to an ongoing outbreak that has great impact on public health and society. Increased use of sequencing technologies in local laboratories can help prevent increase of SARS-CoV-2 spreading by better understanding modes of transmission.

## Introduction

In December 2019 China reported a group of patients with a severe respiratory illness caused by a thus far unknown coronavirus. Severe acute respiratory syndrome coronavirus 2 (SARS-CoV-2) was identified as the causative agent.^1^ Since its outbreak, the virus evolved into a pandemic with almost 37 million infections and over a million deaths worldwide by October 2020.^2^ Many countries are currently fighting second waves of infection whilst the healthcare systems are still under pressure from the first wave. To reduce spread and mitigate workforce depletion, large scale testing of healthcare workers (HCW) was implemented in the Netherlands early on.^3^

Current testing is based on RT-PCR detection of SARS-CoV-2 in nasopharynx or oropharyngeal swabs. If tested SARS-CoV-2 positive, HCW are instructed to self-isolate at home, and source finding and contact tracing is performed. These procedures enable us to identify patients and personnel at risk of infection and to identify chains of transmission in the hospital. In the community setting, source finding and contact tracing is performed by public health staff upon a notification of a SARS-CoV-2 positive individual. It facilitates the implementation of quarantine measures for high-risk contacts in the community. Contact tracing is time consuming and with rising numbers of infections as currently seen in the second wave, the public health capacity may reach the limits of feasibility of thorough source and contact tracing investigations.^4^ Routine sequencing the SARS-CoV-2 genome from positive samples provides crucial insights into viral evolution and supports outbreak analysis.^5,6^ Current whole-genome sequencing (WGS) workflows often require cumbersome preparation, are laborious to implement for high throughput screening or use less widely accessible sequencing platforms, preventing widespread implementation. Here we present a novel strategy for fast, simple and robust Next-Generation Sequencing (NGS) WGS library preparation. We show that the RC-PCR method, which integrates tiled target amplification with Illumina library preparation has a simple workflow with minimal hands-on time. We used this novel and practical method to I) validate and compare it with another sequence technology to demonstrate its reliability and capacity and II) apply it to a set of epidemiologically linked cases to illustrate its added value in detecting potential transmission events in public health and hospital settings.

## Material and Methods

In this study we conducted a validation to assess the performance and reproducibility of the novel RC-PCR SARS-CoV-2 sequencing technology. Subsequently, we performed a clinical validation to assess the potential added value in identifying chains of transmission in a hospital and public health setting.

### Sample collection

Nasopharyngeal and oropharyngeal swabs collected in UTM or GLY medium of patients, healthcare workers and samples for the local public health services that were tested for SARS-CoV-2 in our laboratory. Samples collected between March 2020 and September 2020 were included in this study and stored at −80°C. Detailed descriptions on included samples can be found in supplementary table 1. A total of 173 SARS-CoV-2 positive and fifteen SARS-CoV-2 negative samples were tested.

### Samples and selection of epidemiological clusters

Nineteen out of 188 samples were previously sequenced using Oxford Nanopore Technologies (ONT). These nineteen samples were collected at the beginning of the pandemic, between March 9^th^ and March 20^th^ and ONT sequencing data of these samples has been deposited at GISAID, a global initiative curating sequenced SARS-CoV-2 genomes for public access (https://www.gisaid.org/).^6^

#### Hospital samples

Six epidemiolocal hospital clusters that were identified by the infection prevention and control (IPC) team were included in this study. These clusters involved patients admitted at and healthcare workers (HCW) employed by the Radboud university medical center. Of the identified clusters, three were clusters of healthcare workers with an epidemiological link, and three involved a patient and several healthcare workers with a suspected epidemiological link. To determine whether other HCW could be linked to one of the clusters, samples of sporadic HCW (all other HCW who tested positive for SARS-CoV-2 in September 2020) were included in the selection, as the second wave of infections in the Netherlands started late August 2020. These consist of Radboud university medical center HCW and the majority work in direct or indirect patient care. A minority of positive samples include employees working at the medical faculty or research departments. Additionally, twenty samples were included from patients and HCW who were tested between March and September 2020 and who were not associated with any of these predefined clusters.

#### Community samples

We also included an additional 64 community samples that tested positive for SARS-CoV-2 in March and April 2020 and that were tested by the local public health service. These were samples of persons living in the defined public health region surrounding our hospital. See Table 1 for an overview of the groups and clusters.

**Table 1:** number of groups and clusters of samples that were sequenced for SARS-CoV-2.

|  |  |  |  |
| --- | --- | --- | --- |
| **Groups** | **Samples (N)** | **Month SARS-CoV-2 PCR positive** | **IPC cluster information** |
| Oxford Nanopore Technology (ONT) | 19 | March 2020 | None |
| Cluster 1 – External outbreak link | 6 | September 2020 | HCW linked to a known community outbreak and who had either visited the venue or had close contact to people (with positive test) who had visited the venue |
| Cluster 2 – Department C | 5 | September 2020 | All HCW working at the same department in close proximity and who tested positive in the same week. |
| Cluster 3 – Patient ward E | 2^ | September 2020 | A patient and an HCW; the HCW had contact with the patient without adequate personal protective equipment (PPE). |
| Cluster 4 – Patient ward H | 3* | May 2020 | Two HCW and one patient tested positive at the same department in a short time period. An epidemiological link was suspected since the employees came in contact with the patient. |
| Cluster 5 – Laboratory R | 9 | April 2020 | All HCW working at the same department, tested positive in the same week. |
| Cluster 6 – Patient ward S | 6 | September 2020 | One patient and 5 HCW, the HCW tested positive 5 days after being in contact with the positive patient, the event included an unexpected aerosol generating procedure and HCW were not protected with PPE. |
| Sporadic HCW September 2020 | 39 | September 2020 | none |
| Public Health services samples | 64 | March & April 2020 | none |
| Other (patients/employees tested up to September 2020) | 20^ | March – September 2020 | none |
| Negative | 15 | n.a. | n.a. |
| Total | 188 |  |  |

samples was anonymized. Cluster information was provided anonymously by the IPC team.

The Research Ethics Committee of the region Arnhem/Nijmegen reviewed the current study and waived additional ethical approval. All personal data of patients, HCW and public health service

### Real-Time Polymerase Chain Reaction

SARS-CoV-2 RT-PCR was performed on all samples during routine diagnostics. RNA was isolated using Roche COBAS 4800 (Roche Diagnostics Corporation) with a CT/NG extraction kit according to the manufacturers protocol. RT-PCR with primers targeting the envelope (E-gene) was used as described by Corman *et al*. and performed on a LightCycler 480 (Roche Diagnostics Corporation) using Roche Multiplex RNA Virus Mastermix.^7^

### Reverse Complement Polymerase Chain Reaction

For all 188 selected samples, RNA isolation was repeated on the MagnaPure 96 (Roche Diagnostics Corporation) using Small Volume isolate protocol with 200μl of sample and eluting isolated RNA in 50μl. cDNA-synthesis was performed using Multiscribe RT (Applied Biosystems) with 10μl of RNA input (supplementary table 2). Four samples were replicates, RNA was isolated twice and tested in two separate sequencing runs. They were randomly selected for the first run, but were also part of an IPC identified cluster and therefore included in the second run.

Whole genome sequencing (WGS) was performed in 3 independent runs (96 samples each) using the novel EasySeq™ RC-PCR SARS-CoV-2 WGS kit (NimaGen BV, Nijmegen, The Netherlands). Figure 1 and 2 show a detailed description of the technology in which two types of oligo’s are used to start the targeted amplification. The RC-probe and the universal barcoding primer hybridize and start the formation of specific SARS-CoV-2 primers with Unique Dual Index (UDI) and adapter sequences already included. In contrast to other techniques where multiple steps are needed to add sequence adapters and UDI’s. This means a regular PCR-system can be used to produce SARS-CoV-2 specific amplicons ready for sequencing. The kit uses 155 newly designed probes with a tiling strategy previously implemented in the ARTIC protocol.^8^ The probes are divided in two pools, A and B. Pool A contains 78 probes and Pool B contains 77 probes. This strategy requires two separate RC-PCR reactions but ensures there is minimal chance of forming chimeric sequences or other PCR artifacts (See Figure 2). After the PCR, samples of each plate are pooled into an Eppendorf tube, resulting in two tubes, for pool A and B, respectively. These are individually cleaned using AmpliClean™ Magnetic Bead PCR Clean-up Kit (NimaGen, Nijmegen, The Netherlands). Afterwards, quantification using the Qubit double strand DNA (dsDNA) High Sensitivity assay kit on a Qubit 4.0 instrument (Life Technologies) is performed and pool A and B are combined. The amplicon fragment size in the final library will be around 435 bp. Next Generation Sequencing (NGS) was performed on an Illumina MiniSeq® using a Mid Output Kit (2×150-cycles) (Illumina, San Diego, CA, USA) by loading 0.8 pM on the flowcell. The first two runs (Run1 and Run1_new) were conducted to test the performance of the RC-PCR on a large variety of Ct-values (Ct 16 – 41) using the standard protocol provided by NimaGen. For sequencing Run1_new the RC-PCR product from Run1 was re-used and sequenced with the exception that the final sequencing library was created by using a balanced library pooling strategy based on estimated cDNA input (2 ul for Ct<20, 5 ul 20≤Ct<27 or 10ul Ct≥27). The final sequence run (Run2) contains samples with a Ct range from 16 – 32, using the same Ct dependent balanced library strategy.

**Figure 1.**
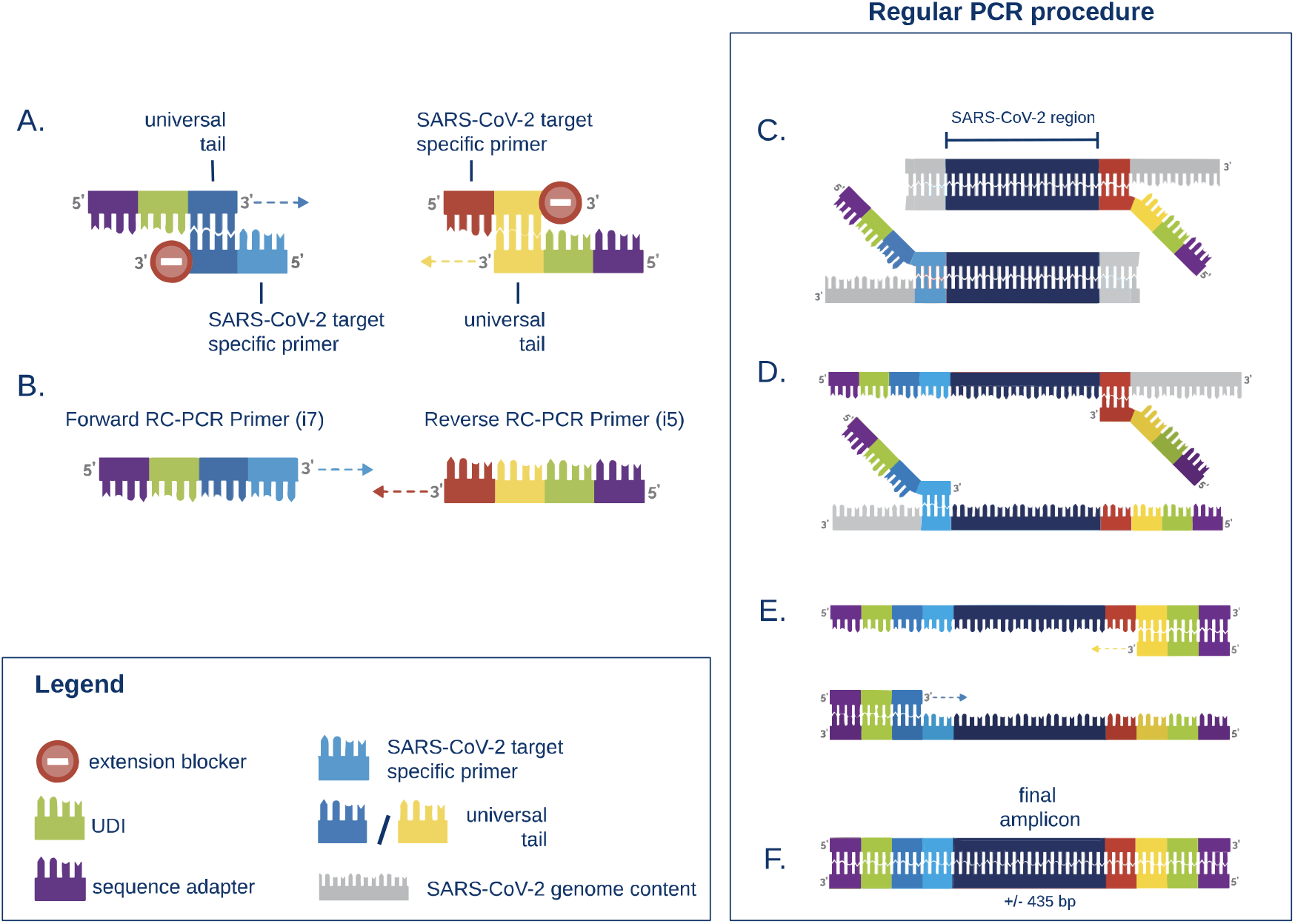
Schematic representation of the RC-PCR technology to WGS SARS-CoV-2. The protocol consists of one single PCR-like reaction consisting of 2 steps. The schematic is adapted from Kieser *et al*. (Kieser *et al*, 2020) **A.** Two types of oligo’s are present, 1) the universal barcoding primer which includes a Unique Dual Index (UDI), sequence adapter, and universal tail. 2) the RC probe which contains an extension blocker, universal sequence, and the reverse complement of the SARS-CoV-2 genomic target sequence. **B.** The universal tail sequences anneal and form a SARS-CoV-2 specific PCR primer. **C - E.** A regular PCR in which the SARS-CoV-2 specific amplicons are created. **F.** The final amplicons are ready to sequence on an Illumina sequencer.

**Figure 2.**
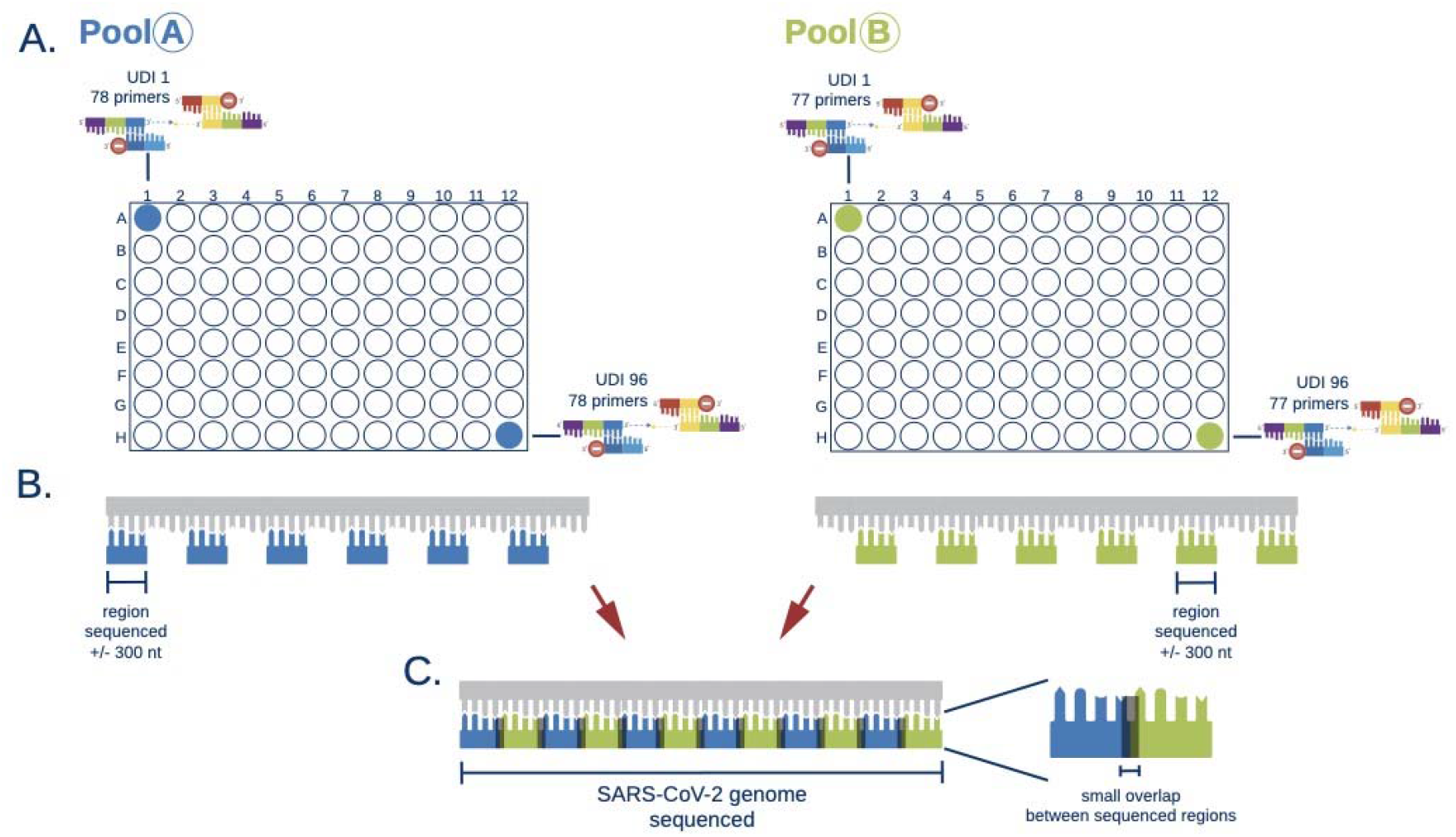
Schematic overview of the EasySeq SARS-CoV-2 WGS process. **A.** For 96 samples the kit provides a Pool A plate and a Pool B plate. These plates contain the RC-probe and the universal oligo’s with an UDI per well. Identical UDI’s between Pool A and B are used. **B.** For covering the full SARS-CoV-2 genome a tiling method similar to the ARTIC protocol is used.(DNA Pipelines R&D, 2020) The kit provides two distinct plates to separate into amplicon pools that do not overlap. This greatly enhances accurate sequencing output. **C.** The same wells of pool A and B share the same index and allow the combination of corresponding sequencing results to cover die full SARS-CoV-2 genome.

### Data analysis

VirSEAK (JSI, Ettenheim, Germany) was used to map the Illumina paired-end reads to SARS-CoV-2 reference NC_045512.2. Consensus sequences were extracted for each sample using the virSEAK export option, settings used can be found in supplementary table 3. All consensus sequences and reference NC_045512.2 were aligned using MUSCLE (version 3.8.1551) using default settings.^9^ Sequence statistics were calculated using faCount (version 377). Mean read depth (RD) was calculated using JSI/SEQUENCE PILOT (JSI, Ettenheim, Germany) to evaluate the amplicon depth of each of the 155 amplicons. For the validation samples (ONT group Table 1) the sequence starts and ends were trimmed to match RC-PCR region with Oxford Nanopore region. A maximum-likelihood phylogenetic tree was inferred using IQ-TREE (version 2.0.3) under the GTR□+□F□+□I□+□G4 model with the ultrafast bootstrap option set to 1,000. Phylogenetic tree visualization and annotation was performed using iTOL (version 5.6.3) or FigTree (version 1.4.4) (http://tree.bio.ed.ac.uk/software/figtree/).^10^ SNP distances between samples was calculated using snp-dists (version 0.7.0) (https://github.com/tseemann/snp-dists). From the genome alignments we calculated a minimum spanning tree (MST) by applying the MSTreeV2 algorithm using GrapeTree (version 1.5.0).^11^ Visualization of the MST was performed using GrapeTree.

The clinical validation consisted of a comparison of the epidemiological information of the community and hospital samples and the WGS findings to see whether sequencing confirmed or dismissed the suspected links between the samples.

## Results

### Technical results RC-PCR

In this study we performed three Illumina MiniSeq Mid Output (2×150 bp) runs containing 96 samples each that were prepared using the EasySeq™ RC-PCR SARS-CoV-2 WGS kit. It has a turnaround time of about 8.5 hours, consisting of 1-hour hands-on time for preparing 96 samples, 6.5 hours for performing the RC-PCR, and 1-hour of hands-on time for pooling, sample clean-up. Run 2 had the highest number of positive SARS-CoV-2 VirSEAK consensus retrievals (100%). Of Run1 65% was retrieved, Run1_new 67%. Run2, containing samples with higher viral loads (Ct values 16-32), reached an average coverage of 96.69%. Genome coverage for Run1_new was 88%. (Figure 3B) Supplementary table 4 provides a detailed overview of the technical results of the three sequence runs.

**Figure 3.**
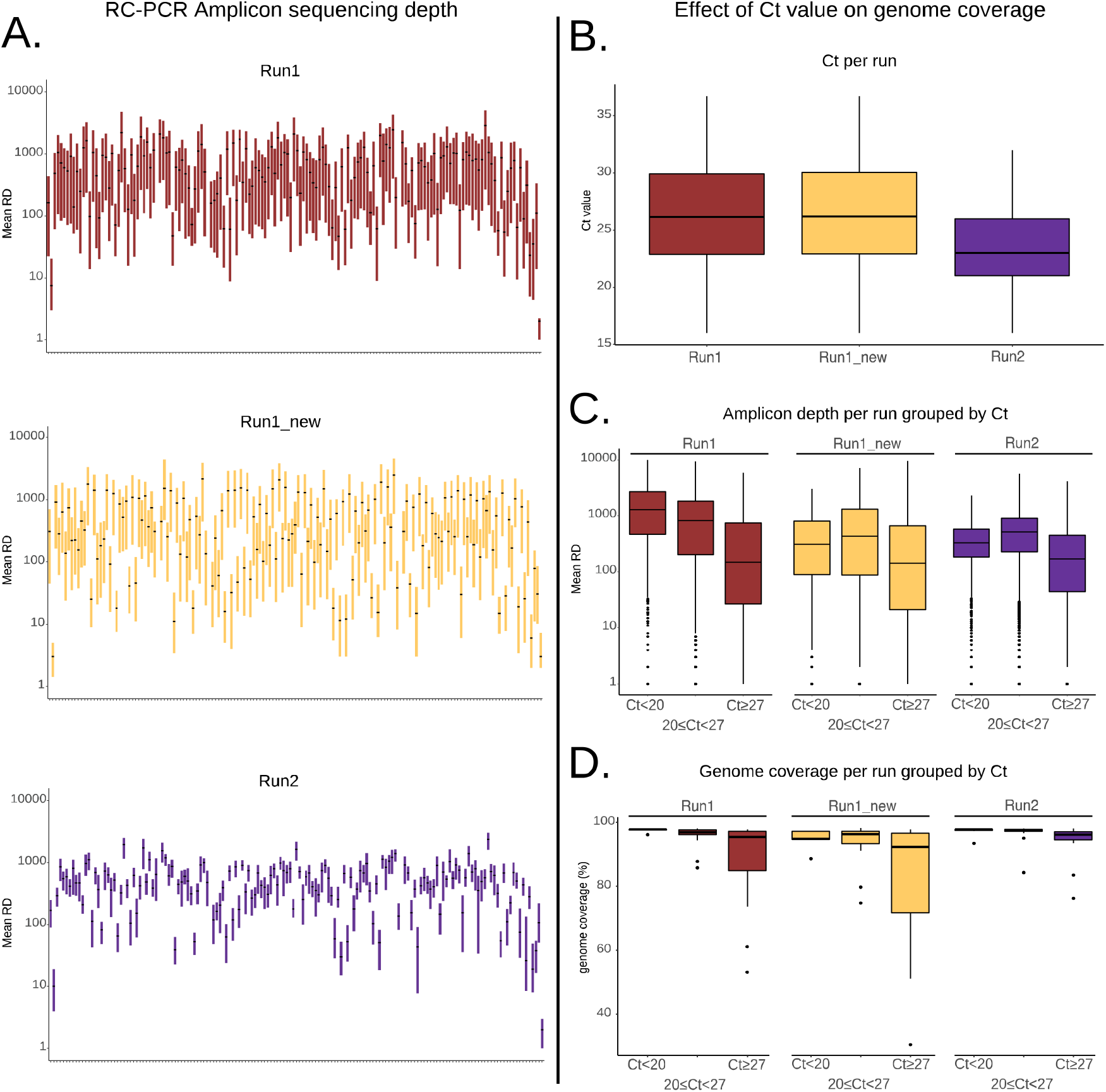
Graphical representation of performance of the RC-PCR Illumina sequence runs. **A.** Boxplots of the interquartile range of the Mean read depth (RD) of the Amplicons on a log_10_ scale for all 155 probes sorted on the SARS-CoV-2 genome. **B.** Boxplot of the Ct value as determined by RT-PCR to illustrate the differences of viral load of the sample per run. **C.** Boxplots of the Mean RD of the amplicons (log_10_ scale used) grouped per run and by Ct value. **D.** Boxplots of the SARS-CoV-2 genome coverage as achieved by RC-PCR grouped per run and Ct value.

#### Amplicon depth plots

The amplicon depth distribution highlights which parts of the SARS-CoV-2 genome are represented and the number of reads for each of the amplicons. In essence this shows how well the individual parts of the SARS-CoV-2 genome are represented in the results. To illustrate the amplicon distribution on the SARS-CoV-2 genome, for each of the 155 amplicons a sequencing depth was calculated and plotted per run (See Figure 3A). Most amplicons are centered around a Mean read depth (RD) of 100-1000. While some amplicons show less depth, in most cases they still result in a consensus sequence. Additionally, for Run2 the interquartile range of the Mean RD is smaller compared to the two other runs. When comparing the amplicon depth obtained per probe, boxplots are made for each run divided in three Ct groups (Ct<20, 20≤Ct<27, and Ct≥27) (see Figure 3C). We see a decline in depth for samples with Ct above 27. For Run1_new and Run2 samples with a Ct between 20 and 27 perform slightly better than the Ct<20 group this is probably an effect of the balanced library input strategy applied for these runs. To evaluate if the impact of amplicon sequencing depth affects SARS-CoV-2 genome completeness, boxplots with the Ct groups are displayed to the effect on genome coverage (see Figure 3D). Here we notice a decline in genome coverage with increasing Ct values for Run1 and Run1_new. Run2 maintains high genome coverages, however does not contain samples with Ct values above 32.

#### Regions of low sequencing coverage

In a detailed analysis of the coverage of the SARS-CoV-2 genome obtained by RC-PCR 5 missing genomic regions were observed (Table 2). The largest missing region has a length of 186 bp and is part of the Open Reading Frame 1a (ORF1a). A further two regions are the start (1-54 bp) and the end (46-165bp) of the genome. We observed that region 14585-14725 is missing in the VirSEAK consensus output but not in the JSI/SEQUENCE PILOT and at the time of writing the manuscript the VirSEAK algorithm was updated to improve the consensus output. Overall, without this update, the maximum SARS-CoV-2 genome coverage that can be achieved using RC-PCR is between 97,8% and 98,2%. In version 1 of the EasySeq™ RC-PCR SARS-CoV-2 WGS kit three probe pairs do not produce amplicons, 6258_6426, 9504_9752, and 21241_21420, respectively. No data on these genomic regions will be obtained (Table 2).

**Table 2:**
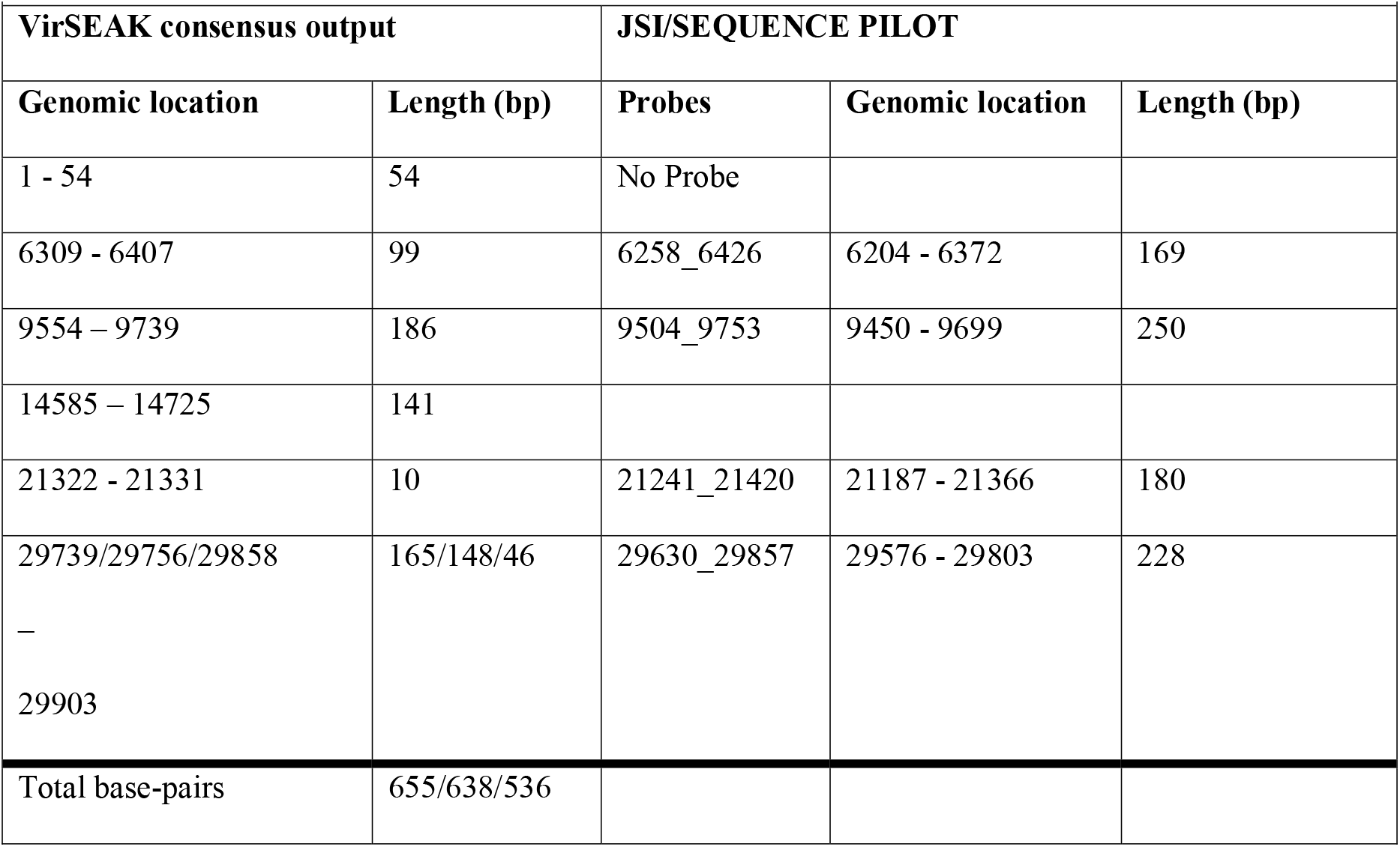
missing regions in VirSEAK consensus output.

### Validation of RC-PCR reproducibility

All samples from Run1 that obtained a consensus (n=57) were compared to the same 57 samples from Run1_new to determine whether results are reproducible when repeating sequencing with the RC-PCR product. Results in supplemental figure 1 show that 50 of the 57 clusters fully align between Run1 and Run1_new. There are 7 samples in which the phylogenetic distance is larger. For those samples in which the phylogenetic distance is larger than expected, alignments were analyzed. The samples from Run1_new show a lower genome coverage, explaining larger phylogenetic distances in these cases.

This is in line with the results observed in table 3 with average genome coverage of 88% in Run1_new versus 93% in Run1. Which is either caused by RC-PCR product storage or the influence of the balanced library pooling strategy based on Ct values of the samples.

Four sample pairs were tested in both Run1 and Run2 to serve as biological replicates. The entire process from RNA isolation to sequence analysis was performed twice on these four samples. Phylogenetic analysis depicted in Figure 4 (Illumina biological replicates) shows perfect agreement between these repeats and confirms the specificity and reproducibility of RC-PCR.

**Figure 4.**
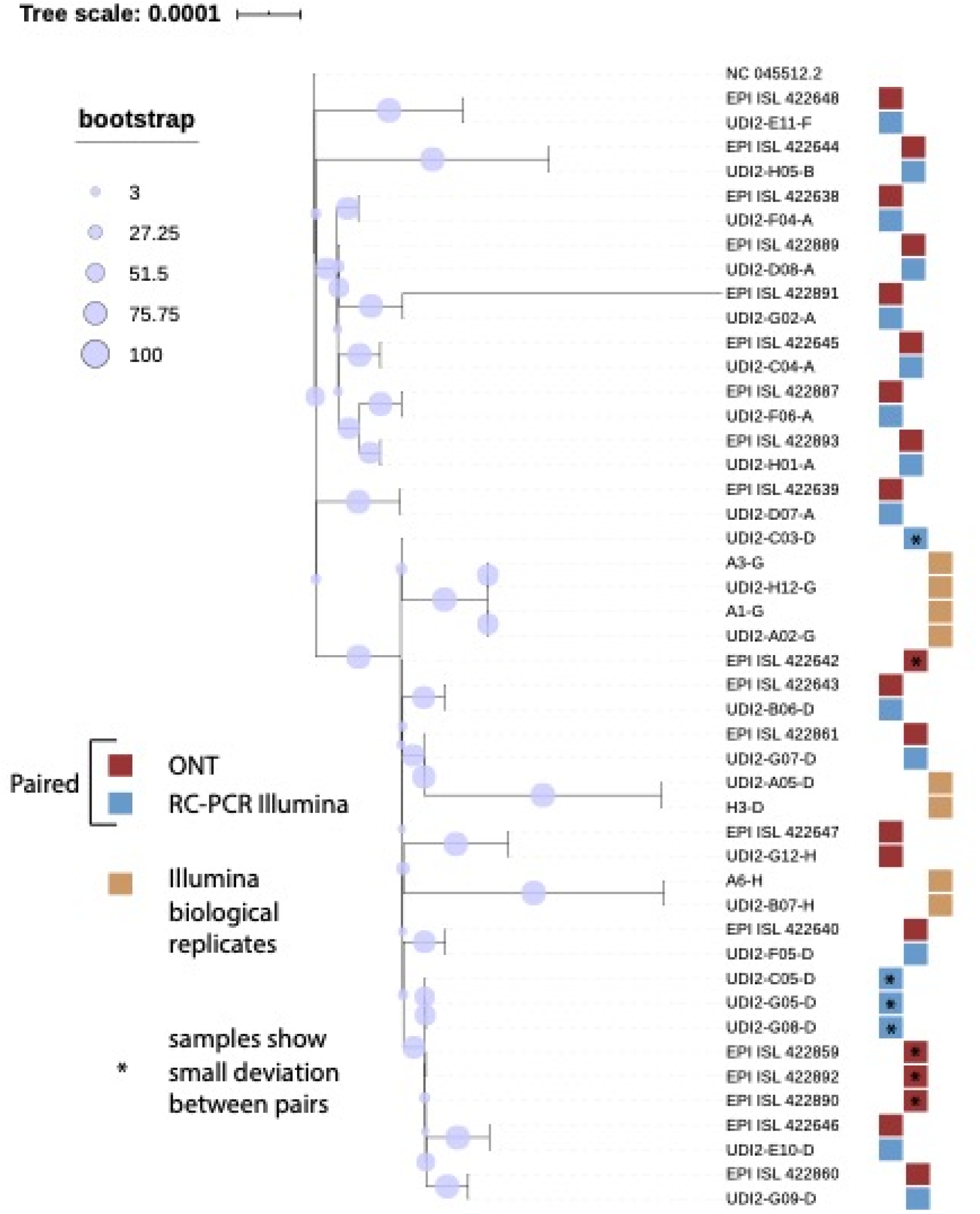
Validation of RC-PCR by comparing the results for the same 19 samples sequenced previously by ONT. (Munnink *et al*, 2020) Additionally, reproducibility was tested by applying RC-PCR. on 4 samples as biological teplicates (beige).

### Validation of RC-PCR with Oxford Nanopore Technologies® (ONT)

Nineteen out of the 188 samples were tested using both ONT and Illumina® sequencing. The ONT sequences were available in the GISAID database and compared to the results of RC-PCR sequencing. All nineteen samples provided sequencing results on both platforms (Figure 4, ONT in red and RC-PCR in blue). Fourteen out of nineteen samples provided perfect pairs, four samples show a small divergence in the phylogenetic tree. Single nucleotide polymorphism (SNP) distance was calculated to identify the number of nucleotides discrepant between samples. This in combination with manual inspection showed that they have identical sequences but RC-PCR samples miss certain genomic regions compared to ONT which results in the phylogenetic differences. One pair does not match, the ONT sample shows a large distance (EPI ISL 422891). Manual inspection of the alignment revealed a wrongly placed ambiguous region in the ONT sample.

### Clinical validation

Of the 188 tested samples, 173 were SARS-CoV-2 positive of which sequencing results were obtained for 146 (57 in Run1 and 89 in Run2). All samples, excluding nineteen ONT and four duplicate samples used for validation, are depicted in the phylogenetic tree of Figure 5. Only HCW and patients are included in the minimum spanning tree of Figure 6. Figure 5 shows the genetic diversity of the samples at different time points during the pandemic. Those collected during the first months of March and April (community samples from public health service) are clearly separated from the other samples, especially compared to the samples from September 2020 (Cluster 1,2,3,6, and the HCW). In Figure 6 it is clear the epidemiological link between the samples three of the six clusters was completely confirmed by the sequencing results. Clusters two, five, and six contained HCW that were not related. In cluster one, linked to a venue outside the hospital, five samples group together with no SNP distances, one sample has a distance of a single SNP suggesting the possibility of linked cases. However, multiple “sporadic HCW” tested in September and two HCW previously linked to cluster two and five also group within cluster one.

**Figure 5.**
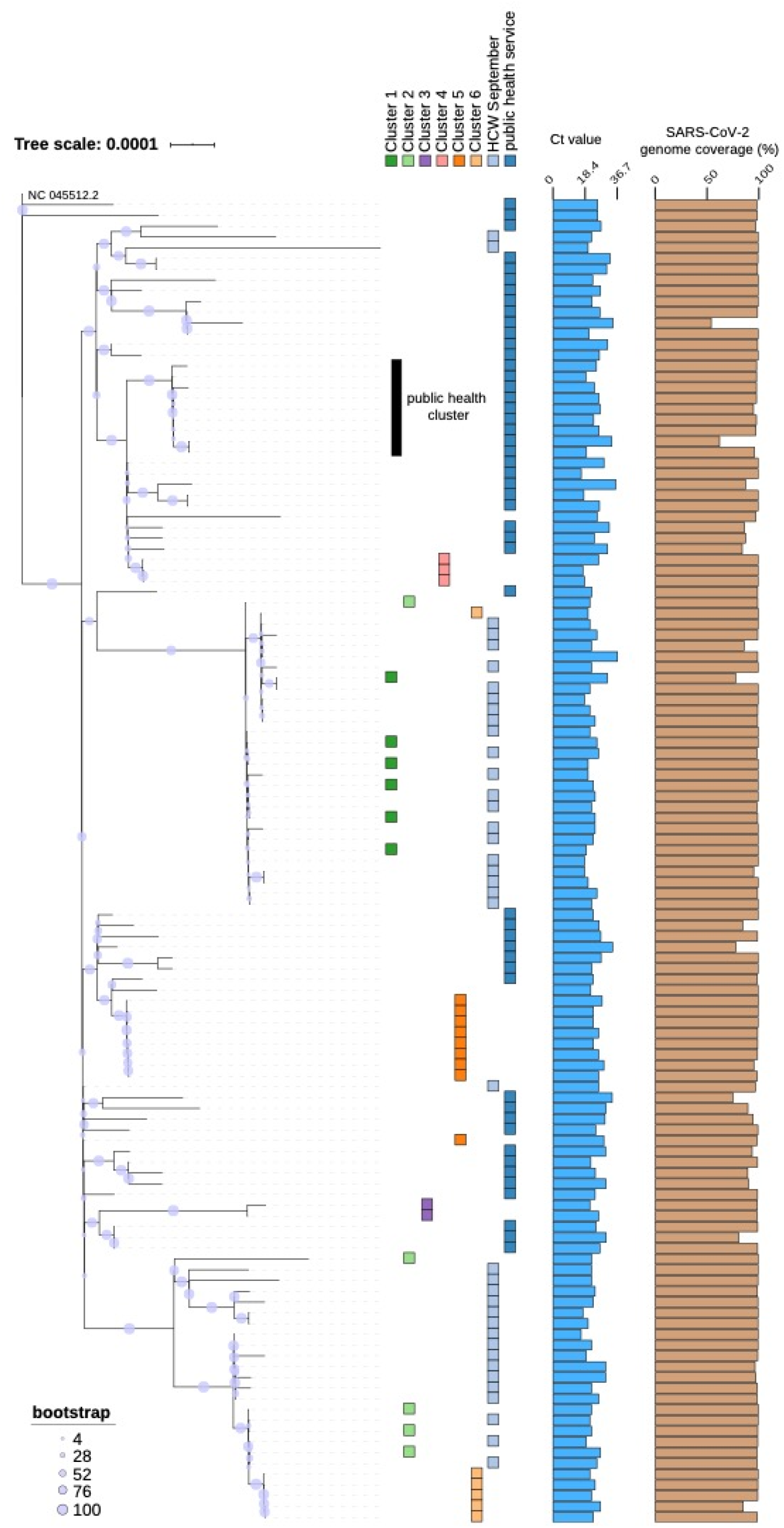
Phylogenetic tree of 123 RC-PCR WGS SARS-CoV-2 genomes rooted at the NC 045512.2 reference genome. These samples are obtained from SARS-CoV-2 positive tested patients, HCW, and samples provided by public health services. Six clusters of samples were identified by the hospital Infection prevention control team. Sample groups are indicated by the colored blocks. Additional Ct values and genome coverage are plotted in barplots to illustrate the diversity in viral load between the samples and the high genome coverage that can be achieved by RC-PCR, respectively.

**Figure 6.**
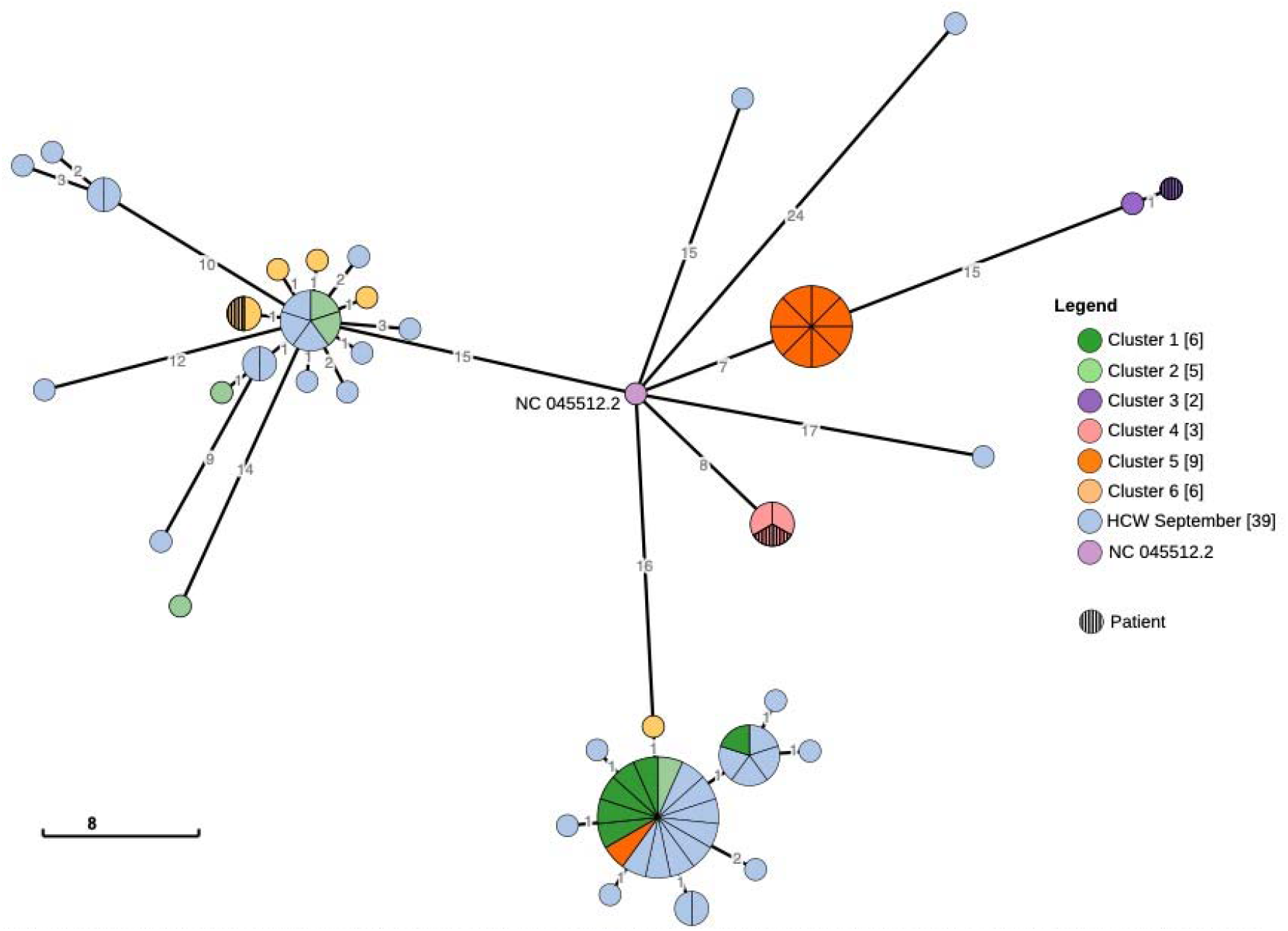
Minimum spanning tree of all samples being part of a cluster as defined by the infection prevention team and all HCW September samples. The tree was calculated using GrapeTree with the MSTreeV2 algorithm. This figure clearly illustrates the relationship between the samples and the clusters.

In cluster two only two samples group together, two others are genetically unrelated samples and one samples has a SNP distance of 2 which could still be within the transmission chain. Cluster three, a patient and HCW show a distance of only 1 SNP. Sample collection was performed on one occasion, twelve days apart, which could account for the SNP difference. In cluster four two HCW and a patient group together, confirming the suspected link. Cluster five, an outbreak at a laboratory, eight HCW samples have identical SARS-CoV-2 genomes, only one sample is phylogenetically linked. Cluster six originated from a SARS-CoV-2 positive patient seen at a department, where at that time multiple HCW had close contact to the patient. At the time of presentation, no symptoms were present that were indicative of SARS-CoV-2 and screening using a questionnaire was negative. Five of the HCW tested SARS-CoV-2 positive in the following weeks. In four HCW a genetically similar SARS-CoV-2 virus was detected. Surprisingly multiple other HCW group in this same cluster with minimal differences (0-3 SNPs), which could mean the outbreak was larger than anticipated or the patient was not the source of the infection.

Even though no new clusters were identified among the “sporadic HCW”, they do group with previously identified clusters. Additional information about these HCW revealed that many of them had a direct or indirect link to the community source that was known by the public health services, Cluster one.

Sequencing of the 64 community samples showed seven people clustered together in the phylogenetic tree of Figure 5. There was no prior information available on these tested persons, but additional information provided by the Local Public Health Service indicated that two of the seven worked at the same location, two were their partners, the others lived in the same neighbourhood at the initial four people, although they had no known epidemiological link to these people other than the area of residence. Of other public health service samples no contact tracing information was available and other samples clustering could not be confirmed with an epidemiological link.

## Discussion

In this study we present the application of a novel method called Reverse Complement-PCR to sequence the SARS-CoV-2 genome which combines target amplification and indexing in a single procedure, directly creating a sequencing ready Illumina library. We applied this method to 173 hospital and community samples that tested positive for SARS-CoV-2 with RT-PCR. Most epidemiological clusters from the hospital and the community were confirmed by phylogenetic clustering. Based on our data, RC-PCR is a reproducible technology, it correlates well with Oxford Nanopore sequencing, is able to sequence samples with Ct values up to 32 determined by RT-PCR and within these samples retrieves a high SARS-CoV-2 genome coverage. Optimization of the protocols is expected to increase coverage in samples with lower viral loads even further.

Previous studies showed the benefit of using WGS of SARS-CoV-2 for outbreak investigation purposes and to study transmission routes.^6,12–16^ Several methods have been optimized for this purpose. The ARTIC Illumina method, a tiling multiplex PCR approach, was the first that enabled WGS of SARS-CoV-2 using Illumina sequencers.^17^ The technique has subsequently been optimized and analysis, albeit in small sample numbers, concluded that it delivers sufficient quality to perform phylogenetic analysis.^18–20^ It had been used as targeted and random RT-PCR screening with subsequent sequencing of the population in order to study the spread through the community.^12^ More recently Sikkema *et al*. were the first to describe the use of SARS-CoV-2 sequencing in healthcare associated infections and identify multiple introductions into Dutch hospitals through community-acquired infections.^5^

SARS-CoV-2 has an estimated mutation rate of 1.12□× □ 10^−3^ substitutions per site per year, which results in 2.8 mutations every month.^21^ The minimum spanning tree of Figure 6 shows several samples with a genetic distance of only a single SNP. With the mutation rate in mind, it is unclear how to relate these clusters since extensive contact tracing information is lacking and interpretation on SNP regarding outbreak management is unknown. Since community samples of September were unavailable, we are unable to determine whether the genetic diversity in the community was low resulting in genetically similar SARS-CoV-2 strains in a hospital setting. However, since sequencing of samples in March and April 2020 clearly resulted in a larger diversity of SARS-CoV-2, and this was early on in the pandemic, it seems more likely that a common source of infection, in- or outside the hospital is the cause. Further research is needed to determine the accepted SNP distance for the use in outbreak analysis.^22^ Although we know minimum spanning trees are often used in outbreak analysis.^5^ It is a simplification of the phylogeny which could result in erroneous conclusions in outbreak analysis. Care should be taken in interpreting these results.

It should be noted that some of the amplicons result in lower coverage than others (See Figure 3). Currently, developments are under way in which a better distribution of the amplicon depth will be achieved resulting in genome coverage that could increase to almost 100%. The difference in genome coverage between Run1 and Run1_new is most likely caused by storage of the library and subsequent pooling on the basis of Ct value of the individual samples, nonetheless, repeated testing at higher Ct values will be needed to confirm this.

With current increase in infections in many countries including the Netherlands and additional measures being put in place to reduce SARS-CoV-2 spreading, real-time sequencing of public health service samples could be used to target infection prevention measures nationwide and locally.^23^ Its application can range from incidental cluster analysis in the case of uncertain epidemiological links to real-time surveillance in the community or health care institutes. Additionally, correlation between specific SARS-CoV-2 strains or mutations and clinical outcome could be identified, supporting clinical decision making to improve outcomes for patients.^24,25^

In conclusion, here we implemented for the first time, RC-PCR in the field of medical microbiology and infectious diseases thereby showing it to be a robust method which requires only minimal hands-on time compared to current sequencing methods and can be used for high throughput sequencing of SARS-CoV-2. Moreover, RC-PCR and sequence analysis can support epidemiological data with genomic data to identify, monitor, and screen clusters of samples to help identify chains of transmission of SARS-CoV-2, enabling a rapid, targeted and adaptive response to an ongoing outbreak that has great impact on public health and society.

## Author contributions

F.W. and J.P.M.C. conducted the research, performed analysis, wrote manuscript and created the figures. L.F.J.vG., C.P.B-R., E.C.T.H.T., N.vdG-B., J.L.A.H. proofreading and provided clinical information and samples of patients and HCWs. A.T.and J.H. conducted the contact tracing and proofreading of the manuscript. H.F.L.W., J.C.R-L., M.S., and W.J.G.M supervised the study and drafted the manuscript.

