## Supplemental data for "Novel SARS-CoV-2 Whole-genome sequencing technique using Reverse Complement PCR enables easy, fast and accurate outbreak analysis in hospital and community settings"

| Supplemental table 1: Sample details of 188 samples (4 duplicates) | | | |
|---|---|---|---|
| **Run 1** | **Sample plate position** | **Cluster** | **Ct-value** |
| Run 1 | A01 | Patient ward H | 17 |
| Run 1 | A02 | Patient June | 34 |
| Run 1 | A03 | Patient ward H | 26 |
| Run 1 | A04 | Patient July | 25 |
| Run 1 | A05 | Employee may | 32 |
| Run 1 | A06 | Patient ward E | 23 |
| Run 1 | A07 | Public health service | 31 |
| Run 1 | A08 | Public health service | 28 |
| Run 1 | A09 | Public health service | 34 |
| Run 1 | A10 | Public health service | 21 |
| Run 1 | A11 | Public health service | 22 |
| Run 1 | A12 | Public health service | 32 |
| Run 1 | B01 | Public health service | 31 |
| Run 1 | B02 | Public health service | 27 |
| Run 1 | B03 | Public health service | 33 |
| Run 1 | B04 | Public health service | 23 |
| Run 1 | B05 | Public health service | 30 |
| Run 1 | B06 | Public health service | 30 |
| Run 1 | B07 | Public health service | 34 |
| Run 1 | B08 | Public health service | 24 |
| Run 1 | B09 | Public health service | 25 |
| Run 1 | B10 | Public health service | 34 |
| Run 1 | B11 | Public health service | 25 |
| Run 1 | B12 | Patient may | 33 |
| Run 1 | C01 | Public health service | 30 |
| Run 1 | C02 | Public health service | 34 |
| Run 1 | C03 | Public health service | 26 |
| Run 1 | C04 | Public health service | 32 |
| Run 1 | C05 | Public health service | 33 |
| Run 1 | C06 | Public health service | 16 |
| Run 1 | C07 | Public health service | 22 |
| Run 1 | C08 | Public health service | 25 |
| Run 1 | C09 | Public health service | 26 |
| Run 1 | C10 | Public health service | 23 |
| Run 1 | C11 | Public health service | 22 |
| Run 1 | C12 | Public health service | 30 |
| Run 1 | D01 | Public health service | 30 |
| Run 1 | D02 | Public health service | 26 |
| Run 1 | D03 | Public health service | 29 |
| Run 1 | D04 | Public health service | 22 |
| Run 1 | D05 | Public health service | 30 |
| Run 1 | D06 | Public health service | 24 |
| Run 1 | D07 | Public health service | 19 |
| Run 1 | D08 | Public health service | 27 |
| Run 1 | D09 | Public health service | 33 |
| Run 1 | D10 | Public health service | 23 |
| Run 1 | D11 | Public health service | 30 |
| Run 1 | D12 | Public health service | 27 |

| | | | |
|---|---|---|---|
| Run 1 | E01 | Public health service | 34 |
| Run 1 | E02 | Public health service | 27 |
| Run 1 | E03 | Public health service | 27 |
| Run 1 | E04 | Public health service | 34 |
| Run 1 | E05 | Public health service | 25 |
| Run 1 | E06 | Public health service | 24 |
| Run 1 | E07 | Public health service | 26 |
| Run 1 | E08 | Public health service | 27 |
| Run 1 | E09 | Public health service | 19 |
| Run 1 | E10 | Public health service | 24 |
| Run 1 | E11 | Public health service | 27 |
| Run 1 | E12 | Public health service | 22 |
| Run 1 | F01 | Public health service | 18 |
| Run 1 | F02 | Public health service | 26 |
| Run 1 | F03 | Public health service | 26 |
| Run 1 | F04 | Public health service | 24 |
| Run 1 | F05 | Public health service | 31 |
| Run 1 | F06 | Public health service | 29 |
| Run 1 | F07 | Public health service | 23 |
| Run 1 | F08 | Public health service | 20 |
| Run 1 | F09 | Patient April | 36 |
| Run 1 | F10 | Patient April | 36 |
| Run 1 | F11 | Patient April | 36 |
| Run 1 | F12 | Patient April | 36 |
| Run 1 | G01 | Patient May | 35 |
| Run 1 | G02 | Patient May | 41 |
| Run 1 | G03 | Employee May | 42 |
| Run 1 | G04 | Patient May | 36 |
| Run 1 | G05 | Patient June | 36 |
| Run 1 | G06 | Public health service | 36 |
| Run 1 | G07 | Employee May | 35 |
| Run 1 | G08 | Patient July | 39 |
| Run 1 | G09 | Patient August | 39 |
| Run 1 | G10 | Patient August | 37 |
| Run 1 | G11 | Public health service | 35 |
| Run 1 | G12 | Public health service | 35 |
| Run 1 | H01 | Public health service | 37 |
| Run 1 | H02 | Patient may | 37 |
| Run 1 | H03 | Employee September | 16 |
| Run 1 | H04 | Employee September | 28 |
| Run 1 | H05 | Employee | neg |
| Run 1 | H06 | Employee | neg |
| Run 1 | H07 | Employee | neg |
| Run 1 | H08 | Employee | neg |
| Run 1 | H09 | Employee | neg |
| Run 1 | H10 | Employee | neg |
| Run 1 | H11 | Employee | neg |
| Run 1 | H12 | Employee | neg |
| Run 2 | A01 | Employee September | 16 |
| Run 2 | A02 | Patient ward H | 17 |
| Run 2 | A03 | Employee September | 17 |
| Run 2 | A04 | Patient ward H | 18 |

| | | | |
|---|---|---|---|
| Run 2 | A05 | Employee September | 18 |
| Run 2 | A06 | Employee September | 18 |
| Run 2 | A07 | Employee September | 18 |
| Run 2 | A08 | External outbreak link | 19 |
| Run 2 | A09 | Employee September | 19 |
| Run 2 | A10 | Employee September | 19 |
| Run 2 | A11 | Patient ward S | 20 |
| Run 2 | A12 | External outbreak link | 20 |
| Run 2 | B01 | Employee September | 20 |
| Run 2 | B02 | Employee September | 20 |
| Run 2 | B03 | Negative | neg |
| Run 2 | B04 | Employee September | 20 |
| Run 2 | B05 | Employee September | 20 |
| Run 2 | B06 | ONT samples | 21 |
| Run 2 | B07 | Patient ward E | 21 |
| Run 2 | B08 | Department C | 21 |
| Run 2 | B09 | Patient ward S | 21 |
| Run 2 | B10 | Employee September | 21 |
| Run 2 | B11 | Employee September | 21 |
| Run 2 | B12 | Employee September | 21 |
| Run 2 | C01 | Employee September | 21 |
| Run 2 | C02 | Employee September | 21 |
| Run 2 | C03 | ONT samples | 22 |
| Run 2 | C04 | ONT samples | 22 |
| Run 2 | C05 | ONT samples | 22 |
| Run 2 | C06 | Department C | 22 |
| Run 2 | C07 | Department C | 22 |
| Run 2 | C08 | Patient ward S | 22 |
| Run 2 | C09 | Department C | 22 |
| Run 2 | C10 | Employee September | 22 |
| Run 2 | C11 | Employee September | 22 |
| Run 2 | C12 | Employee September | 22 |
| Run 2 | D01 | Employee September | 22 |
| Run 2 | D02 | Employee September | 22 |
| Run 2 | D03 | Employee September | 22 |
| Run 2 | D04 | Employee September | 22 |
| Run 2 | D05 | Employee September | 22 |
| Run 2 | D06 | Employee September | 22 |
| Run 2 | D07 | ONT samples | 23 |
| Run 2 | D08 | ONT samples | 23 |
| Run 2 | D09 | Laboratory R | 23 |
| Run 2 | D10 | Laboratory R | 23 |
| Run 2 | D11 | External outbreak link | 23 |
| Run 2 | D12 | Laboratory R | 23 |
| Run 2 | E01 | Patient ward S | 23 |
| Run 2 | E02 | Employee September | 23 |
| Run 2 | E03 | Employee September | 23 |
| Run 2 | E04 | Patient ward S | 24 |
| Run 2 | E05 | External outbreak link | 24 |
| Run 2 | E06 | Employee September | 24 |
| Run 2 | E07 | Employee September | 24 |
| Run 2 | E08 | Employee September | 24 |

| | | | |
|---|---|---|---|
| Run 2 | E09 | Employee September | 24 |
| Run 2 | E10 | ONT samples | 25 |
| Run 2 | E11 | ONT samples | 25 |
| Run 2 | E12 | External outbreak link | 25 |
| Run 2 | F01 | Employee September | 25 |
| Run 2 | F02 | Employee September | 25 |
| Run 2 | F03 | Employee September | 25 |
| Run 2 | F04 | ONT samples | 26 |
| Run 2 | F05 | ONT samples | 26 |
| Run 2 | F06 | ONT samples | 26 |
| Run 2 | F07 | Laboratory R | 26 |
| Run 2 | F08 | Laboratory R | 26 |
| Run 2 | F09 | Laboratory R | 26 |
| Run 2 | F10 | Employee September | 26 |
| Run 2 | F11 | Employee September | 26 |
| Run 2 | F12 | Employee September | 26 |
| Run 2 | G01 | Patient ward E | 26 |
| Run 2 | G02 | ONT samples | 27 |
| Run 2 | G03 | Department C | 27 |
| Run 2 | G04 | Patient ward S | 27 |
| Run 2 | G05 | ONT samples | 28 |
| Run 2 | G06 | Laboratory R | 28 |
| Run 2 | G07 | ONT samples | 29 |
| Run 2 | G08 | ONT samples | 29 |
| Run 2 | G09 | ONT samples | 29 |
| Run 2 | G10 | Laboratory R | 29 |
| Run 2 | G11 | Laboratory R | 29 |
| Run 2 | G12 | ONT samples | 30 |
| Run 2 | H01 | ONT samples | 30 |
| Run 2 | H02 | Employee September | 30 |
| Run 2 | H03 | Employee September | 30 |
| Run 2 | H04 | External outbreak link | 31 |
| Run 2 | H05 | ONT samples | 32 |
| Run 2 | H06 | Employee | neg |
| Run 2 | H07 | Employee | neg |
| Run 2 | H08 | Employee | neg |
| Run 2 | H09 | Employee | neg |
| Run 2 | H10 | Employee | neg |
| Run 2 | H11 | Employee | neg |
| Run 2 | H12 | Patient ward H | 26 |

| Supplementary table 2: cDNA synthesis | |
|---|---|
| | **µl** |
| 25mM $MgCl_2$ | 5.50 |
| 2.5 mM dNTP mix | 5.00 |
| 10x RT Buffer | 2.50 |
| RNase Inhibitor (20U/µL) | 0.50 |
| Multiscribe RT (50 U/µl) | 0.62 |
| Random hexamer (50 µM) | 1.25 |
| | **15,37 µl mix + 10 µl RNA** |
| **RT reaction thermocycler:** 10 min – 25°C / 30 min – 37°C / 5 min – 95°C / ∞ – 4°C | |

| Supplementary table 3: VirSEAK settings | |
|---|---|
| **Settings** | |
| Profile | SARS-CoV-2 |
| Randomly sheared reads | no |
| Genome mapping | no |
| **Single / double direction analysis** | |
| Both direction minimum absolute coverage | 3 |
| **Mutations** | |
| Min % coverage | 90% per dir. |
| Min % coverage homozygosity | 95% |
| Homopolymer | on |
| **Quality Score** | |
| Score threshold | 30 |
| Ignore reads thresh. | 40% |
| Score coverage warning | off |
| **Expert Settings** | |
| Genome Set | Haploid |
| Max mismatches | 20% |

| Supplementary table 4: Run statistics and details | | | |
|---|---|---|---|
| | **Run1** | **Run1_new** | **Run2** |
| Protocol | Default | Ct Library pooling strategy | Ct Library pooling strategy |
| Total Reads | 19209544 | 18564880 | 18094162 |
| PF Reads | 18152046 | 17436566 | 17333104 |
| % Reads Identified (PF) | 89.18 | 93.92 | 88.79 |
| Forward Read (% >= Q30) | 97.15 | 96.41 | 97.43 |
| Reverse Read (% >= Q30) | 96.3 | 95.02 | 97.27 |
| Total number of samples | 96 | 96 | 96 |
| Negative samples | 8 (8%) | 8 (8%) | 7 (7%) |
| VirSEAK Consensus | 57 (65%)* | 59 (67%)* | 89 (100%)* |
| Failed samples | 31 (34%)* | 29 (33%)* | 0 (0%)* |
| Average amplicon depth | 877.8 | 657 | 555.7 |
| Average Genome Coverage | 92.75% | 88.38% | 96.69% |

** % is calculated in relation to total number of samples excluding the negative samples since output is not expected for these samples.*

### Supplemental figures

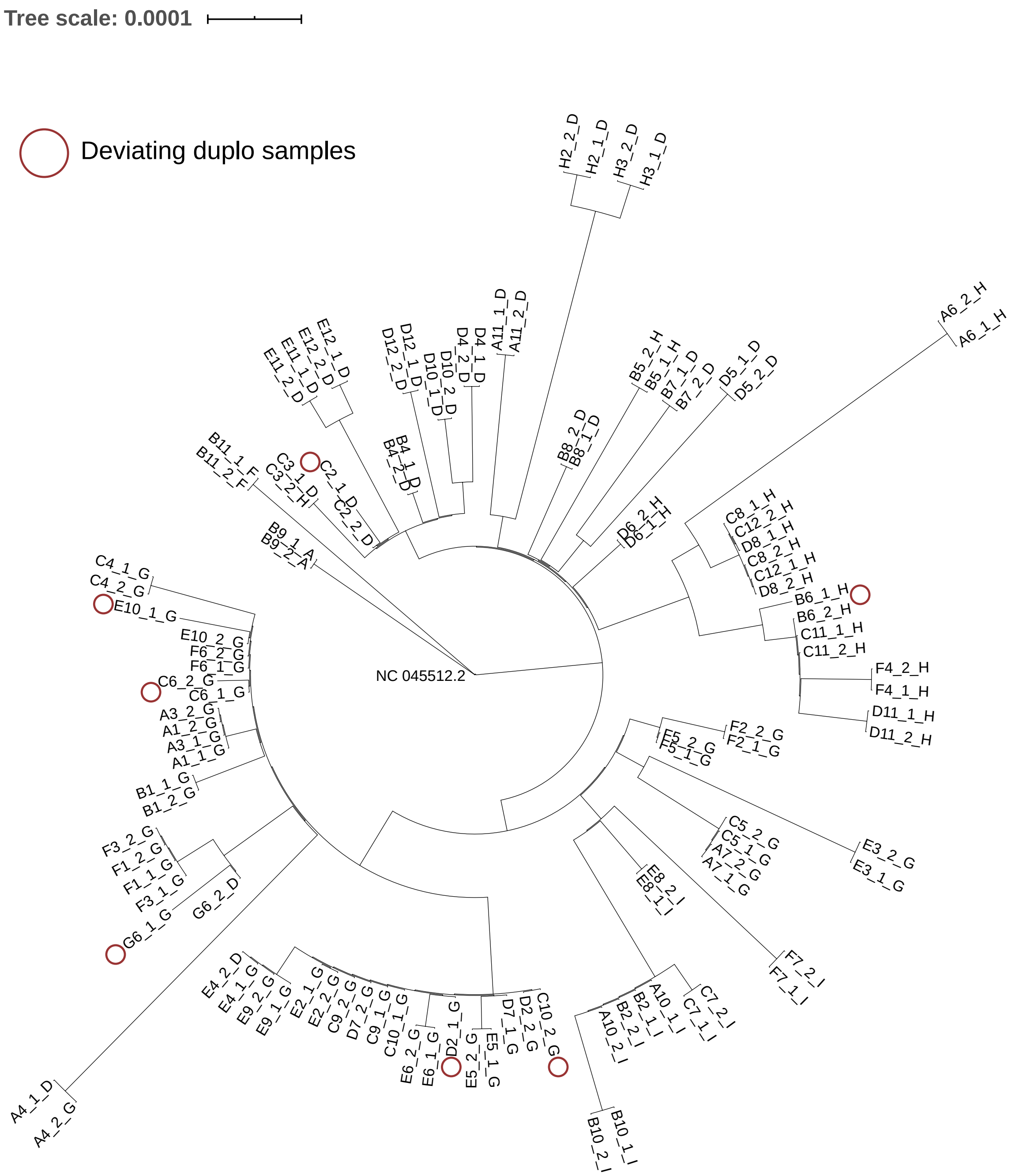


Supplement Figure 1. Validation of RC-PCR by performing a technical duplo of 57 sample of which we performed RC-PCR on the same cDNA. Sample pairs that deviate on the phylogeny are indicated by a red circle.
